# Individual differences in frontoparietal plasticity in humans

**DOI:** 10.1101/2021.11.08.467831

**Authors:** Austin L. Boroshok, Anne T. Park, Panagiotis Fotiadis, Gerardo H. Velasquez, Ursula A. Tooley, Katrina R. Simon, Jasmine C.P. Forde, Lourdes M. Delgado Reyes, M. Dylan Tisdall, Dani S. Bassett, Emily A. Cooper, Allyson P. Mackey

**Author notes:** Austin L. Boroshok (corresponding author) 425 S. University Ave, Stephen A. Levin Bldg., University of Pennsylvania, Philadelphia, PA, 19104. **Author Information** Anne T. Park 425 S. University Ave, Stephen A. Levin Bldg., University of Pennsylvania, Philadelphia, PA, 19104 Panagiotis Fotiadis 240 S. 33rd St., Hayden Hall, University of Pennsylvania, Philadelphia, PA, 19104 Gerardo H. Velasquez 311 Christopher Dr, San Francisco, CA 94131 Ursula A. Tooley 425 S. University Ave, Stephen A. Levin Bldg., University of Pennsylvania, Philadelphia, PA, 19104 Katrina R. Simon 520 W 120th St., Box 199, Russell Hall 21, Teachers College, Columbia University, New York, NY 10027 Jasmine C.P. Forde 11 Mystery Rose Ln, West Grove, PA, 19390 Lourdes M Delgado Reyes 425 S. University Ave, Stephen A. Levin Bldg., University of Pennsylvania, Philadelphia, PA, 19104 M. Dylan Tisdall 3700 Hamilton Walk, Rm. D406, Richards Medical Research Laboratory Bldg. University of Pennsylvania, Philadelphia, PA, 19104 Danielle S. Bassett 240 S. 33rd St., Hayden Hall, University of Pennsylvania, Philadelphia, PA, 19104 Emily A. Cooper 391 Minor Hall, University of California, Berkeley, Berkeley, CA, 94720 Allyson P. Mackey 425 S. University Ave, Stephen A. Levin Building, University of Pennsylvania, Philadelphia, PA, 19104.

## Abstract

Neuroplasticity, defined as the brain’s *potential* to change in response to its environment, has been extensively studied at the cellular and molecular levels. Work in animal models suggests that stimulation to the ventral tegmental area (VTA) enhances plasticity, and that myelination constrains plasticity. Little is known, however, about whether proxy measures of these properties in the human brain are associated with learning. Here, we investigated the plasticity of the frontoparietal system by asking whether VTA resting-state functional connectivity and myelin map values (T1w/T2w ratios) predicted learning after short-term training on the adaptive *n*-back (*n* = 46, ages 18-25). We found that stronger baseline connectivity between VTA and lateral prefrontal cortex predicted greater improvements in accuracy. Lower myelin map values predicted improvements in response times, but not accuracy. Our findings suggest that proxy markers of neural plasticity can predict learning in humans.

## Introduction

Neuroplasticity was canonically defined as the *process* of brain change, but it can also be defined as the brain’s *potential* to change in response to new experiences, and to learn. Animal studies at the level of cells and synapses have made substantial progress in identifying factors that facilitate and constrain neuroplasticity as the potential to change ^1^. Modulatory neurotransmitters, including dopamine, have been shown to increase plasticity ^2^. In a landmark study, stimulation of the ventral tegmental area (VTA), a key source of dopamine for the cortex, restored juvenile-like plasticity in the auditory cortex of adult animals ^3^. Conversely, myelination has been shown to restrict plasticity ^4, 5^. Over 50% of myelin in the cortex is associated with parvalbumin-positive (PV+) inhibitory interneurons ^6, 7^, which are cells that limit synaptic remodeling ^8^. Together, these studies suggest that individual differences in dopamine system connectivity and myelination may contribute to variance in humans’ ability to learn.

The frontoparietal system (FPS) may be a particularly useful target of research on individual differences in plasticity in humans, as the FPS has expanded dramatically over the course of evolution ^9, 10^. The FPS is characterized by a dense expression of dopamine receptors and is lightly myelinated ^11–13^. The FPS is also thought to be highly plastic due to its protracted development ^14, 15^ and high interindividual variability ^16^. High FPS plasticity may be essential for its role as the “multiple demand network” ^17^ and its ability to flexibly adapt its function to meet novel task demands, including working memory and reasoning ^18, 19^.

Some work has been done to understand the process of FPS plasticity: how the FPS changes in response to practice. Long-term reasoning and working memory practice leads to decreases in functional activation in the FPS ^20–27^, and increased functional and structural connectivity between regions of the FPS ^28–32^. FPS can also change over the short-term. Indeed, a few studies have shown that 30-60 minutes of working memory practice is sufficient to cause decreases in FPS activation ^33–35^. In a motor learning task, greater learning was associated with temporal flexibility of functional modules ^36^ and training-related release of coordinated activity across task-extraneous areas ^37^.

However, variability in the FPS’ potential to change is less understood. Investigating variability in the FPS’ potential for change may be important for understanding why some individuals benefit more from educational or cognitive interventions. One study showed that greater gray matter volume in the lateral and medial prefrontal cortex predicted greater learning over five to six weeks of practice with a cognitively complex video game ^38^. Across a broader set of learning tasks, including perceptual and motor learning, greater learning over days or weeks is predicted by greater cortical thickness in task-relevant regions ^39^, greater task activation during the to-be-learned task ^40–42^ and during feedback on a separate task ^43^, and stronger functional connectivity within task-relevant regions ^44, 45^. Long-term training studies, however, are not well-suited to investigating individual differences in the brain’s potential to change. Baseline brain measures may not strongly predict learning outcomes in long-term training studies because variability in learning outcomes over the course of several weeks or months are more likely to be shaped by differences in practice intensity, or by differences in lifestyle factors that influence brain health, including sleep and stress ^46, 47^. It may be better to instead study variability in short-term learning. One study found that positive functional connectivity within the FPS and in other task-positive systems predicted greater learning from 80-90 minutes of working memory practice ^48^. However, these measures are difficult to link to cellular markers of plasticity.

Here, we focused on plasticity as a potential to change rather than plasticity as a process. We examined whether MRI-based proxy measures of FPS plasticity at baseline predicted individual differences in learning following short-term, adaptive working memory training. We identified FPS regions involved in working memory with an *n*-back task, a commonly-used localizer for frontoparietal activation ^49^. As a proxy for dopamine system connectivity, we analyzed resting-state functional connectivity between the VTA and task-active regions. Resting-state functional connectivity is thought to reflect a prior history of coactivation between regions, without confounding effects of task performance ^50^. As a proxy measure for myelin, we examined “myelin map” values, defined as the ratio of T1-weighted (T1w) to T2-weighted (T2w) signal intensities, in task-active regions ^51, 52^. We tested two hypotheses: 1) stronger functional connectivity between the VTA and FPS regions at baseline predicts greater learning following training and 2) lower myelin map values in FPS regions at baseline predict greater learning following training. To evaluate the specificity of the results, we also investigated VTA connectivity and the T1w/T2w ratio in regions we did not expect to be selectively involved in the training task: primary visual cortex and primary motor cortex. Additionally, to explore plasticity as a process, we explored changes in FPS structure and function, and whether these brain changes related to learning.

## Methods

### Ethics Statement

This study was approved by the University of Pennsylvania’s Institutional Review Board. Written informed consent was obtained from all participants.

### Participants

Participants between the ages of 18 and 25 years were recruited through the University of Pennsylvania study recruitment system, as well as through community and university advertisements. Inclusion criteria included fluency in English, no history of psychiatric or neurological disorders or learning disabilities, no current or recent illegal substance use, and no contraindications for MRI.

In total, MRI scans were completed for 61 participants. Forty-six participants were included in the final sample (*M* = 21.39 years, *SD* = 1.91 years; 63% female), which met a predetermined target set by a power analysis indicating that such a sample size would have 80% power to detect a correlation between brain measures and learning of *r* = 0.4. Participants were excluded for falling asleep during the *n*-back scan (*n* = 3), low performance on the control condition of the fMRI *n*-back task (< 90% accuracy on the 1-back condition; *n* = 5), failure to advance beyond the initial working memory condition during the 50-minute training period (*n* = 1), recent illegal substance use not reported during screening but reported during participation (*n* = 1), inability to tolerate scanning (*n* = 1), and technical issues (total *n* = 4; button box malfunction [*n* = 2], coil error [*n* = 1], no behavioral log files [*n* = 1]). The final sample was ethnically and racially diverse (24% Asian, 33% Black, 17% Hispanic/Latino, 4% Multi-Racial, and 19% White; one participant chose not to report their race and ethnicity). 77% of participants were undergraduate students and 18% were graduate students at the University of Pennsylvania.

### Experimental Design and Statistical Analyses

#### Learning measure

Participants completed an auditory *n*-back task outside of the scanner: once before and once after adaptive *n*-back training (Figure 1). The task consisted of four blocks of trials at each of 3 cognitive conditions: 2-, 3- and 4-back (alternating in that order) for a total of 12 blocks. Each of the 12 blocks contained 24 trials. Stimuli were drawn from a pool of eight consonants (‘C’, ‘D’, ‘G’, ‘K’, ‘P’, ‘Q’, ‘T’, and ‘V’). Within each condition, approximately 15% of all trials were targets. At the beginning of each block, the current *n*-back condition was presented in the center of a black screen for 2500 ms, after which the response options “YES” and “NO” appeared. Then, an audio clip of a single consonant played for 500 ms. Participants were given 2000 ms to respond via button press on a standard keyboard: “F” for “YES” responses and “J” for “NO” responses. Each block was followed by 10 seconds of rest. Feedback was provided such that accurate or inaccurate responses prompted the correct response option to be highlighted in green or red, respectively. Two primary indices of learning were analyzed for ease of interpretation: (1) the change in task accuracy across trials, as defined by the percentage change of correctly-answered trials, and (2) the change in response time across trials.

**Figure 1.**
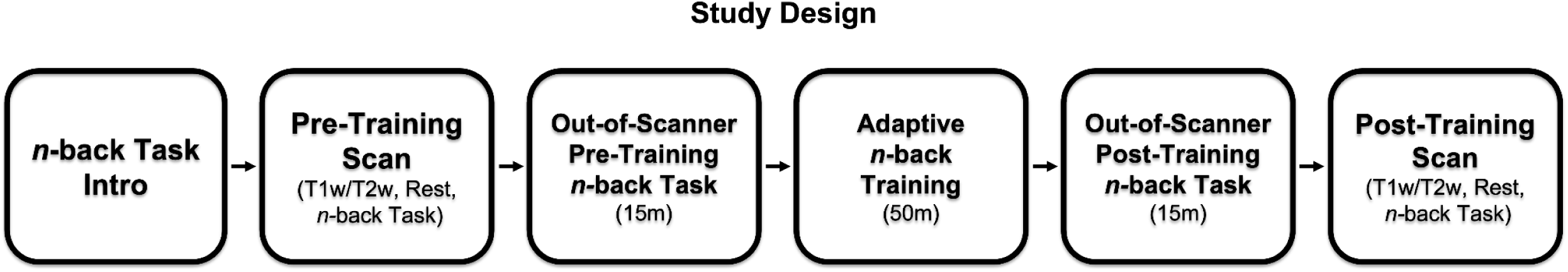
Schematic of study design.

Participants completed a 50-minute adaptive *n*-back task during the training period. The 50-minute duration was selected based on studies that have shown fMRI activation changes following 30-60 minutes of training ^33–35^, and because longer training would likely induce fatigue. The training session was self-paced. Syllables were used during the training period, in contrast to the pre- and post-training assessments, to reduce the likelihood that learning was based on perceptual changes alone. Stimuli were drawn from a pool of eight syllables (‘ba,’ ‘cha,’ ‘da,’ ‘fa,’ ‘ga,’ ‘ja,’ ‘ka,’ and ‘la’). Within each condition, approximately 13% of all trials were targets. The training session began at the 2-back condition; participants progressed to the next-highest task condition if they finished blocks at or above 90% accuracy, remained at the same task condition if they finished blocks with 71-89% accuracy, and regressed to the next-lowest condition (min: 2-back) if they finished blocks with 70% accuracy or below. Each block at each condition consisted of 24 trials. Participants completed a mean of 745 trials (31.1 blocks) and reached, on average, the 5-back condition (min: 3-back, max: 13-back) during the training period.

#### Neuroimaging Data Acquisition

Imaging was performed at the Center for Magnetic Resonance Imaging & Spectroscopy at the University of Pennsylvania with a Siemens MAGNETOM Prisma 3T MRI Scanner (Siemens, Erlangen, Germany) using a 32-channel head coil. Each participant underwent two MRI scans: the first scan was completed before the *n*-back training period and the second scan was completed following training. During both the pre- and post-training scan sessions, participants completed an identical series of scans, which included T1- and T2-weighted structural scans, a resting-state scan, and a five-minute *n-*back task. First, whole-brain, high-resolution, T1-weighted (T1w) multi-echo (MEMPRAGE, TR = 2530 ms; TEs = 1.69, 3.55, 5.41, 7.27 ms; flip angle = 7°; resolution = 1 mm isotropic) and T2-weighted (T2w) structural scans (T2SPACE, TR = 3200 ms; TE = 406 ms; resolution = 1 mm isotropic; turbo factor: 282) were collected with volumetric navigators ^53^. We collected a T2SPACE scan ^52^, but note that this sequence is not a pure T2w scan. The participants viewed a nature documentary during the structural scans. Next, a five-minute run of resting-state fMRI data was acquired (TR = 2000 ms; TEs = 30.20 ms; flip angle = 90°; resolution = 2 mm isotropic). Participants looked at a fixation cross throughout the scan. Resting-state scanning continued until at least 5 minutes of data were acquired with framewise displacement < 0.5 mm. Finally, an *n*-back fMRI scan was acquired (TR = 2000 ms, TE = 30.2 ms, flip angle = 90°, voxel size = 2.0 x 2.0 x 2.0 mm, matrix size = 96 x 96 x 75, 75 axial slices, 170 volumes, field of view = 192 mm). For both EPI sequences, the first four volumes of each scan were automatically discarded to allow time for MRI signal to reach steady-state.

#### fMRI n-Back Task

The fMRI *n-*back task consisted of four 30-second blocks alternating between 1- and 2-back conditions with 12 trials per block. Each block was followed by 10 seconds of rest. Consonant stimuli were presented for 500 ms and participants were given 2000 ms to respond. The fMRI *n*-back task was designed as a FPS localizer; therefore, only 1- and 2-back conditions were used during the fMRI task to maximize accuracy and minimize confounding effects of errors. As a result, the task was not designed to be sensitive to improvements in accuracy associated with training. By design, accuracy on the fMRI *n*-back task was high at baseline (1-back accuracy: *M* = 97.8%, *SD* = 1.9%; 2-back accuracy: *M* = 96.9%, *SD* = 3.5%), and improved from baseline to post-test only during the 2-back condition (1-back: *t* = 0.96, *p* = .34; 2-back: *t* = 2.56, *p* = .01). Response times on the fMRI *n*-back task decreased significantly with training (1-back: *t* = -6.43, *p* < .0001; 2-back: *t* = -3.16, *p* = .002).

### Data Processing and Analysis

#### Task-based fMRI analyses

Preprocessing for the task fMRI data was implemented using FEAT (FMRI Expert Analysis Tool) Version 6.00, part of FSL (FMRIB’s Software Library, www.fmrib.ox.ac.uk/fsl). The following steps were applied: motion correction using MCFLIRT ^54^, skull stripping, spatial smoothing using a Gaussian kernel of FWHM 5 mm, and high-pass temporal filtering (100s). Functional data were normalized to the MNI template during a two-step process using FLIRT (FMRIB’s Linear Image Registration Tool) in FEAT. First, each participant’s functional image was co-registered to their anatomical T1w structural image using FSL’s Boundary-Based Registration feature (BBR) ^55^. Second, the anatomical image was warped to the standard 2 mm MNI152 structural template. Finally, both of these transformations were combined and used to normalize the functional image to standard MNI space ^56^. Average framewise displacement across the pre-training *n-*back run was not significantly different from motion across the post-training run (*t* = .25, *p* = .80). No included participants had average head motion greater than 0.15 mm across either the pre- or post-training runs of the *n*-back task.

At the single-subject level, we created a general linear model (GLM) for each participant that included the following regressors: 1- and 2-back blocks convolved with the double-gamma hemodynamic response function and their temporal derivatives, as well as FSL’s standard and extended motion parameters (global signal, 6 motion parameters and their temporal derivatives, quadratic terms, and the temporal derivatives of the quadratic terms). We conducted mixed-effects analyses in FEAT (FLAME 1) to create group-level maps for three contrasts (1-back greater than baseline; 2-back greater than baseline; 2-back > 1-back) for Pre-training-only, Post-training-only, and Pre-training > Post-training. Examining all three contrasts allowed us to check that the task was activating the regions we expected based on prior work (Supplemental Figure 1). Group-level *z*-statistic images were thresholded using clusters determined by *z* = 4.0 and a corrected cluster significance threshold of *p* = 0.05 ^57^. Results were registered to the Freesurfer *fsaverage* surface and projected to the cortical surface using *mri_vol2surf* for visualization (Freesurfer v6.0) ^58^.

#### Region-of-interest (ROI) definition

To identify task-active ROIs, we used the results of the whole-brain analysis of the 2-back > 1-back contrast from the pre-training scan at a threshold of *z* = 4.0. We selected the five most significant clusters, which were within the frontoparietal system (Figure 2): (1) left lateral prefrontal cortex (left LPFC; MNI coordinates for center-of-gravity: X = -37, Y = 10, Z = 41), (2) right lateral prefrontal cortex (right LPFC; MNI coordinates for center-of-gravity: X = 29, Y = 12, Z = 52), (3) bilateral medial prefrontal cortex (mPFC; MNI coordinates for center-of-gravity: X = -1, Y = 16, Z = 47), (4) bilateral parietal cortex (including medial parietal regions; MNI coordinates for center-of-gravity: X = -4, Y = -56, Z = 48,), and (5) striatum (MNI coordinates for center-of-gravity: X = -7, Y = 3, Z = 12). To examine whether FPS regions specifically predict learning, we also examined two control regions: primary visual cortex and primary motor cortex. We defined these ROIs using the pericalcarine and precentral gyrus regions of the Harvard-Oxford probabilistic cortical structural atlas provided through FSL.

**Figure 2.**
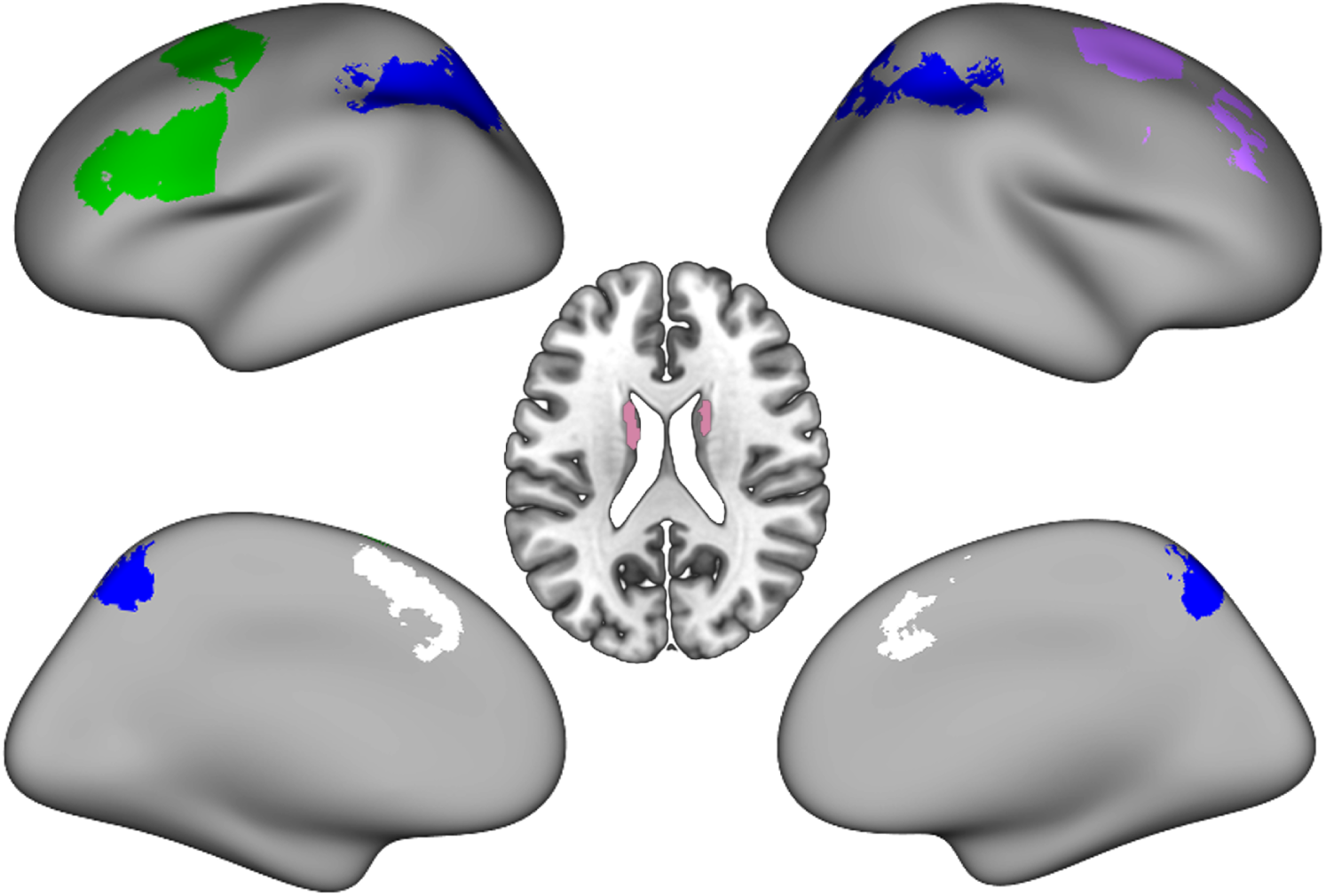
Five task-based regions of interest (left lateral prefrontal cortex [LPFC], green; right LPFC, purple; medial PFC, white; bilateral parietal cortex, blue; striatum, pink) derived from the whole-brain analysis of the 2-back > 1-back contrast from the pre-training scan at a threshold of *z* = 4.0. Axial slice is shown at *z* = 22.

#### Resting-state analyses

Resting-state data were preprocessed with a different pipeline than the one used for the task-based data in order to incorporate Nipype, a Python-based framework specifically optimized for flexibly integrating resting-state analysis tools ^59^. The software packages used in this preprocessing pipeline included FMRIB Software Library (FSL v5.0.8) ^60^, FreeSurfer (v6.0) ^58^, Advanced Normalization Tools (ANTs v2.1.0;) ^61^, and Nipype’s implementation of Artifact Detection Tools (ART; http://www.nitrc.org/projects/artifact_detect/). Simultaneous realignment and slice timing correction were conducted using an algorithm implemented in Nipy ^62^. Outlier volumes in the resting-state data were defined using ART based on composite motion (> 0.5 mm of head displacement between volumes) and global signal intensity (> 3 SD from the mean). Participants had average composite head motion of < 0.2 mm (*M* = 0.11 mm, *SD* = 0.06 mm) across resting-state runs.

The resting-state data were then bandpass filtered (0.01–0.1 Hz), spatially smoothed with an isotropic 6 mm Gaussian kernel (FWHM), and normalized to the OASIS-30 Atropos template (in MNI152 2 mm space) in a two-step process. First, the median functional image was coregistered to the reconstructed surfaces using FreeSurfer’s *bbregister* ^55^; second, the structural image was registered to the OASIS-30 Atropos MNI152 template using ANTs. The transformation matrices generated by these two steps were then concatenated, allowing images to be transformed directly from functional to MNI space in a single interpolation step. The CSF and white matter segmentations were derived from Freesurfer’s individual segmentations of the lateral ventricles and total white matter, respectively, and were transformed into functional space. Five principal components were derived from both segmentations and regressed from the resting-state data, in order to correct for physiological noise like heart rate and respiration (aCompCor) ^63^. At the single-subject level, the following confounds were regressed out: 6 realignment parameters (3 translations, 3 rotations) and their first-order derivatives, outlier volumes flagged by ART (one nuisance regressor per outlier), composite motion, 5 principal components from aCompCor, and linear and quadratic polynomials in order to detrend the data. Global signal was not regressed out during these analyses.

The VTA ROI was defined using a probabilistic atlas ^64^. The average time series of the VTA seed was extracted from unsmoothed functional data and correlated with the average time series from within each of the five task-based ROIs, both before and after training. VTA connectivity with the other ROIs was not related to age, sex, total length of scan (number of volumes collected) or the number of ART outliers (*p*-values > 0.05); nonetheless, we controlled for these measures in all models to ensure that they did not drive relationships with learning. All results of resting-state analyses are the same with and without these covariates of no interest.

#### Myelin maps

We calculated the ratio between each participant’s T1w and T2w images to create subject-specific myelin-enhanced contrast images, or “myelin maps” ^52^, using the publicly available *MRTool* toolbox (version 1.4.3; https://www.nitrc.org/projects/mrtool/) ^51, 65^ for SPM12.

The steps taken by *MRTool* to generate each participant’s myelin map are delineated below. First, each participant’s T2w image was coregistered to their T1w image using a rigid-body transformation ^12, 66^. Both images then underwent bias correction to ensure spatial equalization of the coil sensitivity profiles. The intensity inhomogeneity correction tool in SPM12 was separately used on both images to correct for transmission-field inhomogeneities in image intensity and contrast ^12^. Subsequently, the intensity values of both bias-corrected images were separately standardized using a non-linear external calibration approach (*MRTool* image calibration option #1: Non-linear histogram matching – external calibration), in order to accurately capture inter-individual differences in myelin contrast ^51, 65^. This was a three-step process: (i) subject-specific masks corresponding to CSF, skull, and soft tissues (i.e., dura mater) were extracted using SPM’s Segmentation tool in both anatomical (T1w) and template (MNI) space, (ii) intensity histograms for all three masks were generated in both spaces and a non-linear mapping function (cubic spline interpolation) between them was computed, and (iii) the corresponding cubic polynomial was used to calibrate the intensities of the bias-corrected T1w and T2w images. Lastly, the ratio between each participant’s bias-corrected and calibrated T1w and T2w images was calculated, as a proxy for their corresponding “myelin map” ^52^.

We then masked out CSF and white matter (as defined by individual segmentations in Freesurfer’s LookUp Table) ^67–69^ from the myelin maps to ensure that extracted values for the ROIs reflected only gray matter. The five task-based ROIs (created in MNI space) were inverse-transformed to each subject’s structural space with ANTs ^61^. Finally, myelin map values (T1w/T2w ratio intensities) were extracted from each ROI for each subject, both before and after training.

#### Statistical Analyses

All statistical analyses were performed using *R* (version 4.05) and *RStudio* (version 1.4.1106) software (*R* Foundation for Statistical Computing, Vienna, Austria). We used linear models to predict learning gains (change in accuracy and response time on the n-back task) with resting state functional connectivity (rsFC) between VTA and each ROI, and with T1w/T2w ratios in each ROI. The VTA connectivity models included the following covariates: age, sex, baseline n-back task performance, motion during the baseline resting-state fMRI scan, and the number of volumes acquired during the baseline resting-state fMRI scan. The T1w/T2w ratio models included the following covariates: age, sex, and baseline n-back task performance. All results underwent FDR-correction in R for 28 tests (7 ROIs [5 task-based, 2 control], 2 learning measures [accuracy, response time], and 2 neural measures [VTA rsFC, T1w/T2w ratio].

#### Data Availability Statement

All behavioral data, task-based and control regions of interest (ROIs), values extracted from neuroimaging data, code used to collect functional task data, and code necessary to replicate results are freely available at https://github.com/austinboroshok/frontoparietal-plasticity.

Deidentified neuroimaging data in BIDS format are freely available at https://openneuro.org/datasets/ds003849/versions/1.0.0.

## Results

### Working memory performance improved with training

We considered two behavioral measures of learning: accuracy change and response time (RT) change on the out-of-scanner pre- and post-training *n*-back task (Figure 1). Fifty minutes of training led to small but significant increases in accuracy and decreases in response times (Table 1, Figure 3B). However, there was considerable variability in training gains among individuals (Figure 3A-C). Individuals with lower baseline accuracy improved more on accuracy following training (*β* = -0.284, 95% CI [-0.493, -0.076], *p* = .009). Individuals with slower baseline response times showed swifter response times following training (*β* = -0.389, 95% CI [-0.563, -0.215], p < .001). Gains in accuracy were not associated with improvements in response times (*β* = 0.050, 95% CI [-0.045, 0.145], *p* = .291). Improvements in learning were not significantly associated with age (accuracy: *β* = -0.003, 95% CI [-0.008, 0.002], *p* = .217; RT: *β* = 0.008, 95% CI [-0.008, 0.023], *p* = .306) or sex (accuracy: *β* = 0.019, males higher, 95% CI [0.000, 0.039], *p* = .055; RT: *β* = 0.002, 95% CI [-0.061, 0.065], *p* = .955), controlling for baseline performance. The highest *n*-back condition reached during training was not used in brain analyses because the distribution was significantly non-parametric (Shapiro-Wilk: *W* = 0.81, *p* < .001), due to a small number of participants reaching very high conditions (Figure 3C).

**Figure 3.**
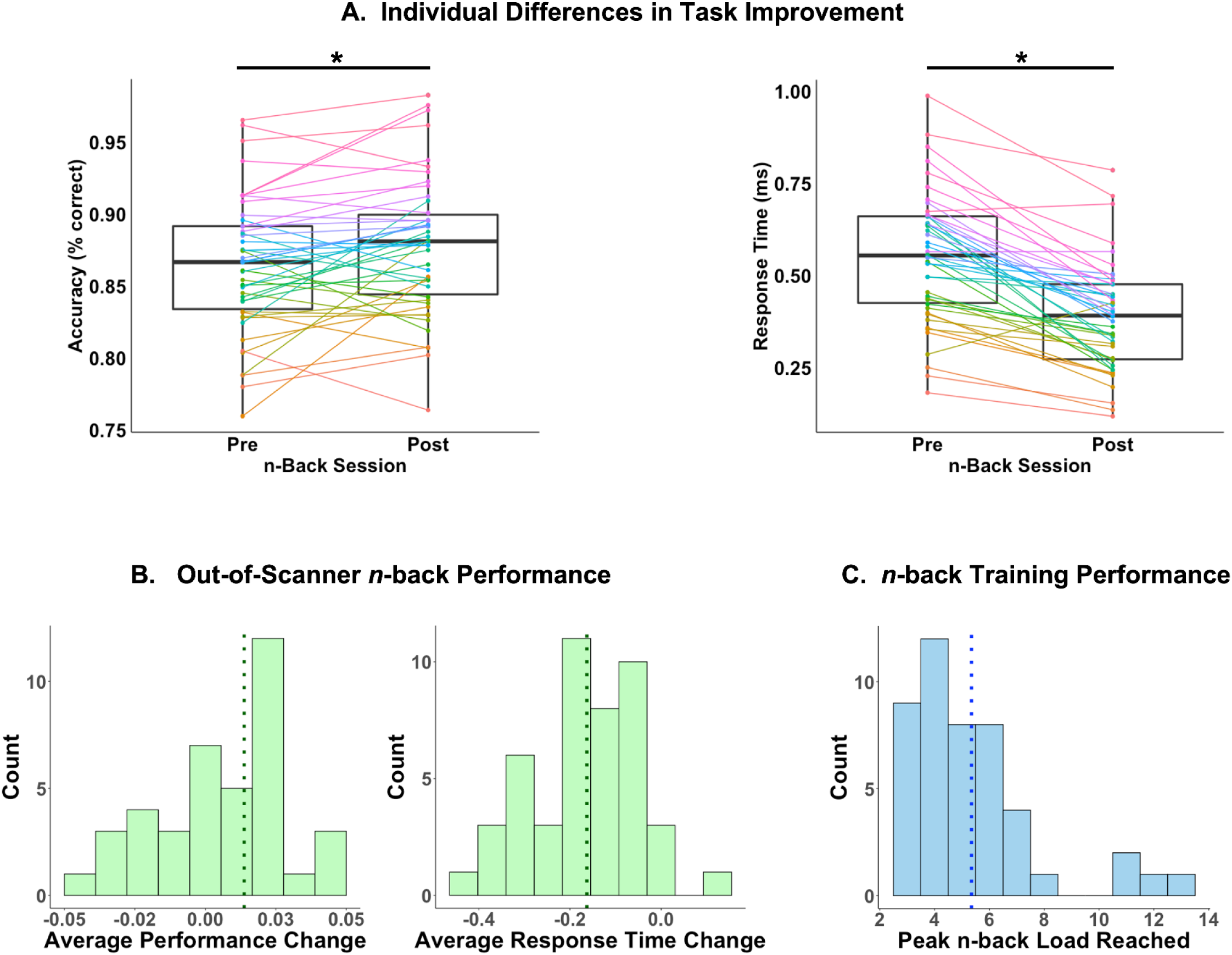
(A) Individual differences in out-of-scanner *n*-back accuracy and response time (averaged across conditions) changes. Each colored line represents an individual participant. (B) Task performance on the out-of-scanner pre- and post-training *n*-back task, measured by accuracy (or percentage of correct trials), averaged across task conditions (left panel) and response time (in seconds) averaged across tasks conditions (right panel). Vertical dashed lines represent mean values. (C) Highest *n*-back condition achieved during training.

**Table 1.** Behavioral performance on the out-of-scanner *n*-back task. Task accuracy is represented using percentage of correctly-answered trials, and response time is represented using seconds taken to respond to stimuli. The delta values for change in accuracy and response time across timepoints represent the difference between pre- and post-training means (*M*) and standard deviations (*SD*). Bolded values denote significant training-related changes, as indicated by Student’s *t*-test. * = *p* ≤ 0.05, ** = *p* ≤ 0.01, *** = *p* ≤ 0.001.

|  | 2-back |  |  | 3-back |  |  | 4-back |  |  | Average |  |  |
| --- | --- | --- | --- | --- | --- | --- | --- | --- | --- | --- | --- | --- |
| | Pre<br>*M*<br>(*SD*) | Post<br>*M*<br>(*SD*) | $\Delta$<br>*M*<br>(*SD*) | Pre<br>*M*<br>(*SD*) | Post<br>*M*<br>(*SD*) | $\Delta$<br>*M*<br>(*SD*) | Pre<br>*M*<br>(*SD*) | Post<br>*M*<br>(*SD*) | $\Delta$<br>*M*<br>(*SD*) | Pre<br>*M*<br>(*SD*) | Post<br>*M*<br>(*SD*) | $\Delta$<br>*M*<br>(*SD*) |
| Accuracy (%) | 92.2<br>(.05) | 94.2<br>(.04) | **.02\*\***<br>**(.04)** | 85.1<br>(.06) | 85.6<br>(.07) | .004<br>(.05) | 82.0<br>(.06) | 83.7<br>(.06) | **.02\***<br>**(.05)** | 86.4<br>(.05) | 87.8<br>(.05) | **.01\*\***<br>**(.03)** |
| Reaction Time (s) | .50<br>(.17) | .39<br>(.17) | **-.11\*\*\***<br>**(.12)** | .60<br>(.21) | .42<br>(.17) | **-.18\*\*\***<br>**(.16)** | .55<br>(.20) | .35<br>(.15) | **-.20\*\*\***<br>**(.17)** | .57<br>(.19) | .41<br>(.16) | **-.16\*\*\***<br>**(.13)** |

### Plasticity as potential

#### Stronger VTA connectivity at baseline predicted greater improvements in accuracy

We used functional connectivity between the VTA and the task-based FPS regions of interest as a proxy measure for dopamine system connectivity (Figure 4A). Consistent with the hypothesis that greater strength of dopamine system connectivity is associated with greater learning, we found that stronger resting-state functional connectivity between the VTA and the bilateral LPFC at baseline predicted greater improvements in accuracy (Figure 4B-C: left LPFC: *β* = 0.067, 95% CI [0.011, 0.123], *p* = .020, *pFDR* = .068; right LPFC: *β* = 0.086, 95% CI [0.027, 0.146], *p* = .006; *pFDR* = .029), controlling for baseline accuracy, age, sex, motion, and total number of volumes. The relationship between VTA-LPFC connectivity and accuracy gains survived FDR correction for 28 tests (7 ROIs, 2 learning measures, and 2 neural measures) for the right LPFC but not the left LPFC. There were no significant associations between accuracy gains and VTA connectivity with the mPFC (*β* = 0.004, 95% CI [-0.057, 0.064], *p* = .906, *pFDR* = .906), the parietal cortex (*β* = 0.019, 95% CI [-0.042, 0.079], *p* = .536, *pFDR* = .578), or the striatum (*β* = 0.050, 95% CI [-0.008, 0.108], *p* = .091, *pFDR* = .213). Further, there were no significant associations between VTA-FPS connectivity and changes in response times in any ROI (left LPFC: *β* = -0.152, 95% CI [-0.329, 0.025], *p* = .090, *pFDR* = .213; right LPFC: *β* = -0.127, 95% CI [-0.322, 0.069], *p* = .198, *pFDR* = .298; mPFC: *β* = -0.119, 95% CI [-0.303, 0.065], *p* = .199, *pFDR* = .298; parietal: *β* = -0.136, 95% CI [-0.316, 0.043], *p* = .133, *pFDR* = .248; striatum: *β* = -0.084, 95% CI [-0.276, 0.108], *p* = .383, *pFDR* = .450). There were no significant associations between VTA connectivity and accuracy or response times at baseline.

**Figure 4.**
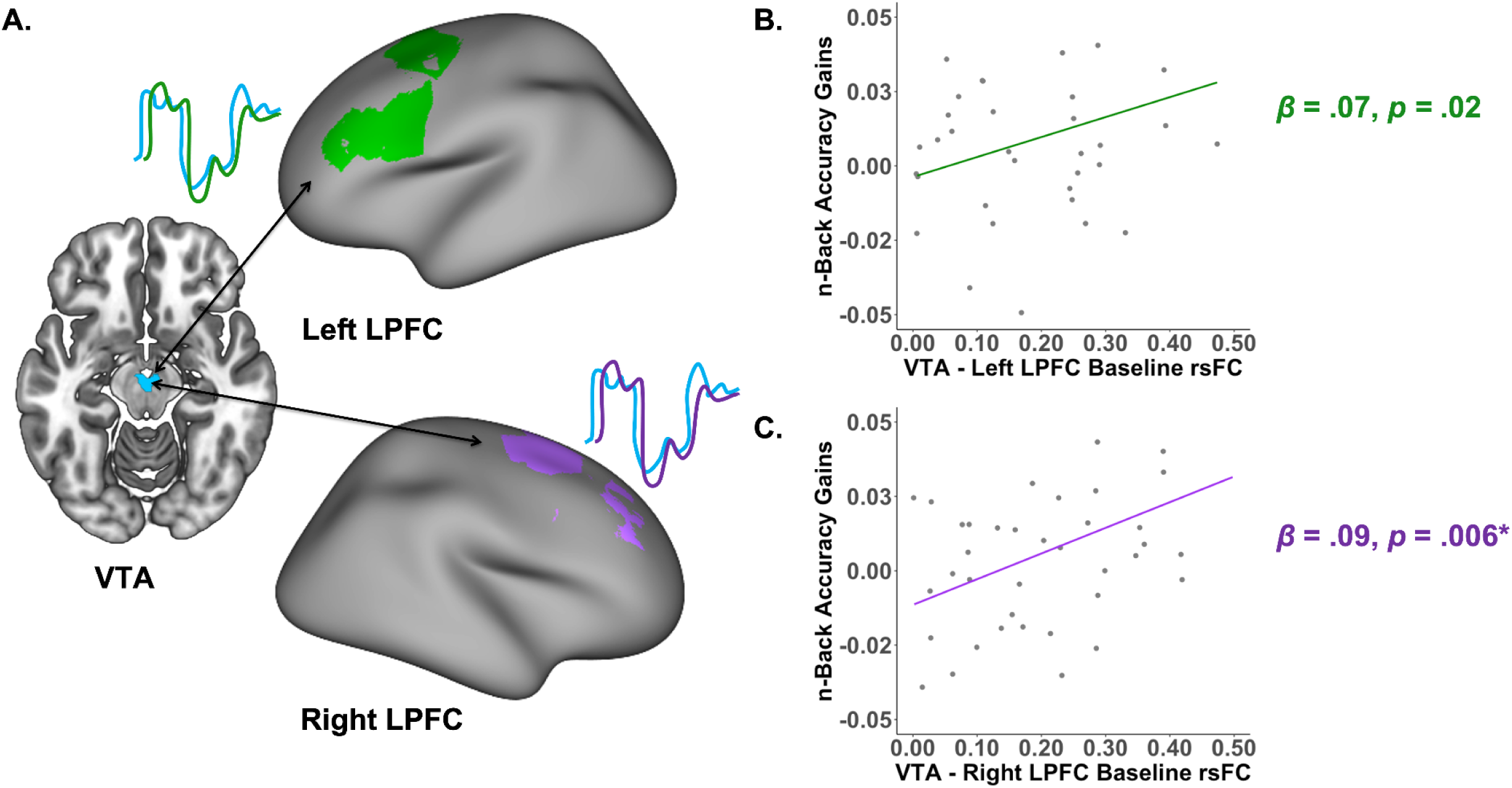
Correlations between functional connectivity between the ventral tegmental area (VTA) and the lateral prefrontal cortex (LPFC) and improvements in accuracy. (A) Schematic of correlation between BOLD time series at rest of regions of interest. Colored lines represent the time series of positive functional connectivity between the VTA and the LPFC at rest. (B) Positive relationships between baseline VTA-left LPFC resting-state functional connectivity (rs-FC) and accuracy gains on the *n*-back working memory task. (C) Positive relationships between baseline VTA-right LPFC rs-FC and accuracy gains on the *n*-back working memory task. Relationships between connectivity and accuracy gains did not survive FDR correction over 28 tests (7 ROIs [5 task-based, 2 control], 2 learning measures [accuracy, response time], and 2 neural measures [VTA rsFC, T1w/T2w ratio]). Statistical models control for age, sex, motion, and baseline working memory accuracy. Asterisks denote *p*-values that survive FDR-correction.

#### Lower T1w/T2w ratios at baseline predicted greater improvements in response times

We used the ratio of T1w/T2w intensities as a proxy measure for myelination (Figure 5A). Individuals with lower baseline T1w/T2w ratios in all five FPS regions of interest showed greater improvements in response times (Figure 5B-F: left LPFC: *β* = 0.423, 95% CI [0.127, 0.719], *p* = .006, *pFDR* = .029; right LPFC: *β* = 0.425, 95% CI [0.131, 0.718], *p* = .006, *pFDR* = .029; mPFC: *β* = 0.318, 95% CI [0.016, 0.620], *p* = .039, *pFDR* = .122; parietal cortex: *β* = 0.410, 95% CI [0.112, 0.708], *p* = .008, *pFDR* = .032; and striatum: *β* = 0.411, 95% CI [0.138, 0.684], *p* = .004, *pFDR* = .029), while controlling for baseline response times, age, and sex. T1w/T2w ratios were not associated with accuracy improvement in any of the task-based ROIs (left LPFC: *β* = 0.075, 95% CI [-0.018, 0.169], *p* = .112, *pFDR* = .223; right LPFC: *β* = 0.075, 95% CI [-0.018, 0.169], *p* = .112, *pFDR* = .223; mPFC: *β* = 0.054, 95% CI [-0.041, 0.149], *p* = .261, *pFDR* = .348; parietal: *β* = 0.068, 95% CI [-0.026, 0.163], *p* = .149, *pFDR* = .261; striatum: *β* = 0.078, 95% CI [-0.009, 0.164], *p* = .077, *pFDR* = .213). Lower baseline T1w/T2w ratios in striatum were related to faster response times at baseline (*β* = 0.491, 95% CI [0.004, 0.979], *p* = .048), while controlling for age and sex. Associations between T1w/T2w ratios and baseline response times in the other FPS regions of interest were not significant (left LPFC: *β* = 0.467, 95% CI [-0.069, 1.002], *p* = .086; right LPFC: *β* = 0.444, 95% CI [-0.092, 0.979], *p* = .102; mPFC: *β* = 0.318, 95% CI [-0.228, 0.863], *p* = .246; parietal cortex: *β* = 0.469, 95% CI [-0.066, 1.004], *p* = .084). T1w-/T2w ratios were not associated with accuracy at baseline.

**Figure 5.**
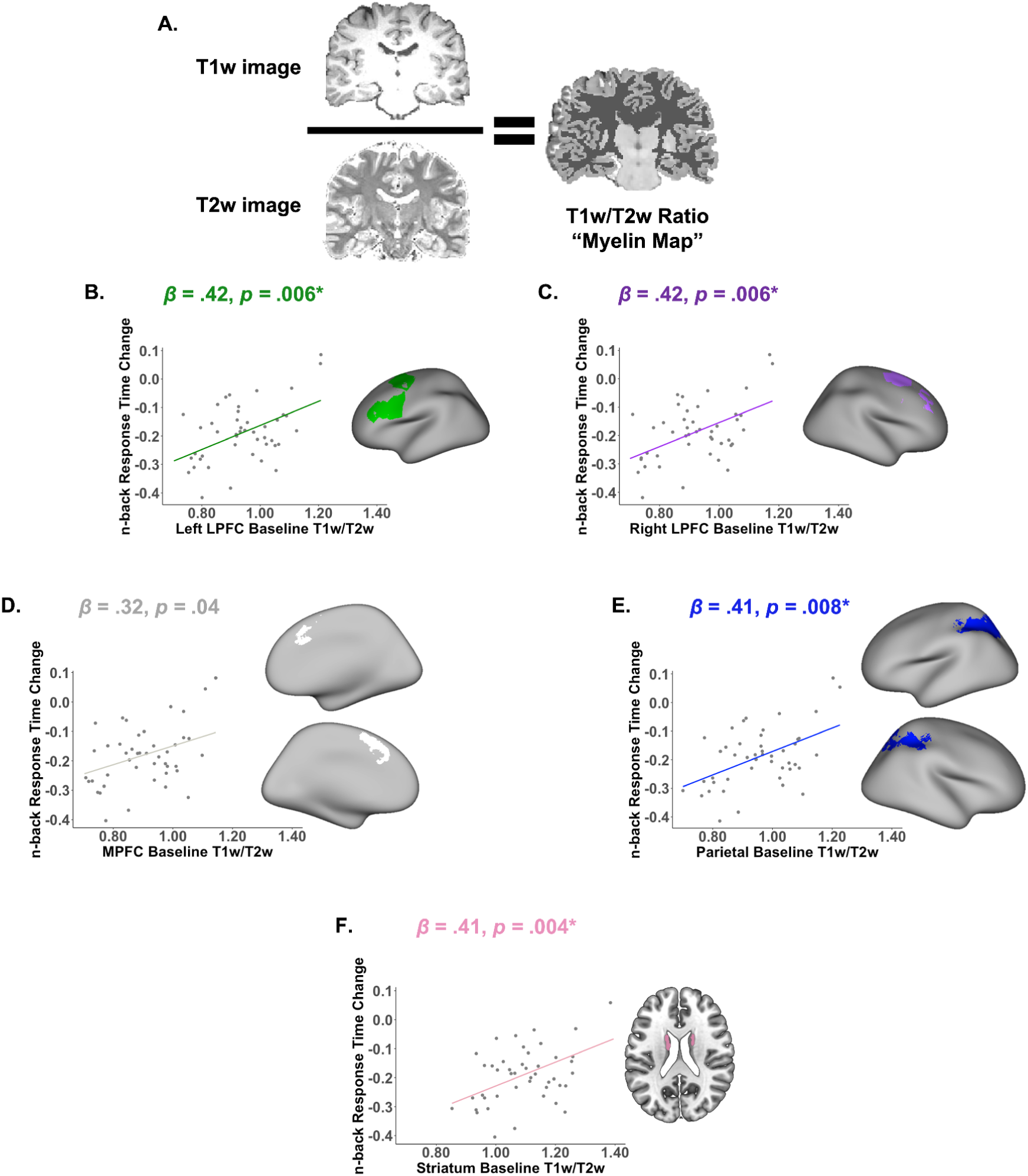
Correlations between frontoparietal T1w/T2w signal intensity ratio and training-related changes in response time. (A) Schematic of T1 signal intensity divided by T2 signal intensity, as a proxy measure for myelination. (B) Task-based regions of interest (B-F) Positive relationships between baseline frontoparietal T1w/T2w signal intensity ratio and training-related changes in response times on the *n*-back working memory task. Axial slice is shown at z = 22. Statistical models control for age, sex, and baseline working memory response times. Asterisks denote *p*-values that survive FDR-correction.

#### Sensitivity analysis

To examine the specificity of the predictions in the frontoparietal system, we examined two control ROIs that we did not expect to predict learning: primary visual and motor cortex. Resting-state functional connectivity between the VTA and visual and motor ROIs did not predict changes in accuracy or response time following training. Baseline T1w/T2w ratios in visual and motor ROIs were not associated with changes in accuracy. However, lower baseline T1w/T2w ratios were associated with greater improvements in response time (visual: *β* = 0.407, 95% CI [0.150, 0.665], *p* = .005, *pFDR* = .01; motor: *β* = 0.426, 95% CI [0.136, 0.715], *p* = .003, *pFDR* = .01).

#### Plasticity as a process: Brain activation changes with training were small and not strongly associated with learning

We did not observe training-related changes in VTA connectivity, or in T1w/T2w ratios (Supplemental Table 1). We also did not observe training-related changes in functional activation (Supplemental Figure 1). Brain changes were not associated with learning. Relationships among brain change measures and learning for all regions of interest are shown in Supplemental Figure 2.

## Discussion

We investigated whether individual differences in the structure and function of the frontoparietal system predicted learning potential in healthy human adults. We focused on proxy measures of two properties that have been shown to influence the brain’s ability to change in animal models: ventral tegmental area (VTA) functional connectivity and myelin maps. Adults, on average, improved their accuracy and response times after 50 minutes of practice on an adaptive *n*-back task, and there were large individual differences in learning. Improvements in accuracy were positively associated with resting-state functional connectivity between the VTA and the bilateral lateral prefrontal cortex (LPFC) at baseline. Improvements in response times were negatively associated with myelin map values for all frontal, parietal, and striatal regions of interest.

The finding that stronger functional connectivity between the VTA and the LPFC predicted greater accuracy gains is consistent with work in animal models showing that dopamine promotes synaptic plasticity ^3, 70, 71^. One interpretation of our findings is that individuals with stronger connectivity between the dopamine system and the LPFC are better able to learn an LPFC-dependent task (i.e., a working memory task) ^72^ because they have greater synaptic plasticity in these regions, or are better able to modulate synaptic plasticity. Indeed, a few experimental and computational modeling studies have suggested that synaptic plasticity in the LPFC is key for working memory ^21, 73–75^. Another interpretation is that individuals with greater top-down control from the LPFC to the VTA are better able to learn because they can better upregulate a range of motivational processes including effort. A third interpretation is that greater VTA-LPFC connectivity reflects a history of coactivation of these regions, perhaps because an individual has more experience learning novel prefrontally-dependent tasks. In rats, the VTA and the PFC show simultaneous and significant increases in firing rate at the same phases of a learning task ^76^. In the present study, it is not the case that VTA-LPFC connectivity is simply a marker of better working memory: VTA-LPFC connectivity was not associated with baseline working memory, and all statistical models predicting working memory change with VTA-LPFC connectivity controlled for baseline working memory. However, resting-state fMRI cannot distinguish between top-down control of the VTA by the LPFC and bottom-up innervation of the LPFC by the VTA, and it also cannot distinguish between dopamine system connectivity and excitatory, or even inhibitory, transmission. Although rs-fMRI leaves open some questions about possible interpretations of our results, VTA connectivity is nevertheless a promising marker of how well adults will learn a frontoparietal task, with potential broader extensions to measuring individual differences in plasticity across other brain systems. Convergent data from PET imaging or pharmacological manipulation of dopamine would strengthen these results.

The observation that individuals with lower T1w-/T2w ratio values in the FPS (a proxy for myelination in this system) at baseline showed the biggest improvements in response time following training is consistent with work in animal models suggesting that lower myelination is associated with greater plasticity ^4, 77^. Less-myelinated individuals were able to improve their response time without sacrificing their accuracy on the *n*-back task, even controlling for baseline performance, alleviating concerns about a speed-accuracy tradeoff effect. Thus, it was not the case that individuals were simply responding faster and less carefully due to increased familiarity with the task or decreased effort following training, which would have been indicated by a high error rate. It remains unclear why our two proxy measures of neuroplasticity were differentially related to the two learning measures. It is possible that the brain measures reflected differences in strategy or approach to the learning task, or that the behavioral outcome measures were sensitive to different learning processes. More work is needed to understand why some individuals show accuracy gains and others show speed gains. We also observed that associations between T1w/T2w ratios and improvements in response time were significant in primary visual and motor regions in addition to our FPS regions of interest. This may indicate that this effect is not specific to the FPS as expected and rather is representative of a more global property of cortex.

Training did not lead to changes in task activation, VTA functional connectivity, or T1w/T2w ratios. Again, as the *n*-back task was designed to serve as a localizer for investigations of FPS *potential* for change, the task was not intended to be particularly sensitive to training-related improvements in accuracy. However, it is possible that changes from short-term learning were too small to be reflected in the neural measures we selected. Additionally, training-related changes in neural measures were not associated with learning gains. It is possible that individuals take different strategies to learn the *n*-back task, and these strategies result in heterogeneous changes in structure and function. Indeed, interindividual variation in learning strategies have been demonstrated to modulate underlying brain structure in a number of learning studies ^78–81^. One such study showed that individual differences in cognitive style and encoding strategies explained significant variability in task-related functional activation during a memory retrieval task in a number of brain regions including frontal and parietal areas ^79^. Another study found similarly strong strategy-dependent changes in lateral prefrontal task-related activation during a working memory task ^81^. Further work in this area is needed to better understand the contributions of individual strategy during short-term learning.

The current study has several limitations. First, the observed relationship between VTA-Right LPFC connectivity and learning survived correction for multiple comparisons, but the relationship between VTA-Left LPFC connectivity and learning did not. Thus, replication of these results is necessary. Second, we conducted analyses that were narrowly tailored to our specific hypotheses about plasticity, but the multimodal data set that we have collected lends itself to additional data-driven analyses, for example asking whether baseline connectivity within the FPS predicts learning ^48^. We share all behavioral and imaging data to facilitate this future work. Third, the study was designed to characterize how brain features predict individual differences in learning, not to test for main effects of working memory training on neural measures, so it did not include a control group. A control group that practiced an unrelated task, for example an implicit learning task that does not rely on the FPS, could further illuminate the specificity of the relationships presented here. For example, a control group could be used to answer the questions: “Does VTA-LPFC connectivity predict accuracy gains on a task that does not engage LPFC?” or “Do T1w/T2w ratios predict swifter response times on a task that does not engage LPFC?” Fourth, we only collected five minutes of resting-state fMRI data from each participant, which may limit the reliability of the VTA-LPFC connectivity findings ^82^. Fifth, our sample included predominantly undergraduate and graduate students, so it may not reflect the variability in cognition and learning that is present in the American population or in the broader world. Finally, learning during the working memory task likely depends not only on the plasticity of the frontal, parietal, and striatal regions, but also on individual differences in effort, attention, strategy choice, or susceptibility to fatigue.

In sum, individuals with stronger connectivity between VTA and lateral prefrontal cortex, as well as individuals with lower myelin map values, showed greater learning from short-term practice. Our study underscores the opportunities and challenges of using neuroimaging tools to measure frontoparietal system plasticity in humans. Better measures of human brain plasticity would enable investigations of the experiences and lifestyle factors that increase plasticity in adulthood, for example stress ^83^, sleep ^84, 85^, or novel positive experiences ^86, 87^. MRI measures of plasticity are also necessary for tackling questions about how early life experiences shape plasticity, with implications ranging from learning in school to response to cognitive behavioral therapies. Therefore, a deeper understanding of human neuroplasticity may help optimize neurocognitive, educational, and psychological interventions that aim to improve well-being and experience across the lifespan.

## Author Contributions Statement

A.L. Boroshok, A.P. Mackey, E.A. Cooper, M. Dylan Tisdall, K.R. Simon, and J.C.P. Forde developed the study concept. All authors contributed to the study design. Testing and data collection were performed by A.L. Boroshok, A.T. Park, G.H. Velasquez, U.A. Tooley, K.R. Simon, and J.C.P. Forde. A.L. Boroshok performed the data analysis and interpretation under the supervision of A.P. Mackey. P. Fotiadis, M. Dylan Tisdall, and E.A. Cooper made substantial contributions to methodological and analytic tools. A.L. Boroshok and A.P. Mackey drafted the manuscript, and A.T. Park, G.H. Velasquez, P. Fotiadis, U.A. Tooley, K.R. Simon, J.C.P. Forde, L.M. Delgado Reyes, M. Dylan Tisdall, D.S. Bassett, and E.A. Cooper provided critical revisions. All authors approved the final version of the manuscript for submission.

## Acknowledgments

We first thank all of the individuals who participated in this research. We also thank Adam Pines, Quentin Wedderburn, Yoojin Hahn, Stephanie Bugden, Ph.D., Ava Cruz, Samantha Ferleger, Destiny Frazier, Jessica George, Abigail Katz, Connor Kendzora, Anna Meaney, Leah Sorcher, Aparna Ramanujam, Daniel Southwick, Alexis Broussard, and Andrea Gomez for their help with data acquisition and cleaning. Finally, we would like to thank Marco Ganzetti for his valuable input towards the usage of the *MRTool* toolbox.

## Supplement

**Supplemental Table 1.** Means and standard deviations of neural measures at baseline and following training, as well as training-related changes in these features. Bold values with an asterisk denote significant training-related changes, as indicated by Student’s *t*-test. * = *p* ≤ 0.05. Abbreviations: lateral prefrontal cortex (LPFC), medial prefrontal cortex (MPFC), resting-state functional connectivity between the ventral tegmental area and frontoparietal regions of interest (VTA rsFC).

|  | *Pre-Training*<br>*M (SD)* | *Post-Training*<br>*M (SD)* | *Change*<br>*M (SD)* |
| --- | --- | --- | --- |
| **VTA rsFC** |  |  |  |
| Left LPFC | .12 (.17) | .08 (.19) | -.04 (.23) |
| Right LPFC | .21 (.16) | .23 (.17) | .02 (.21) |
| MPFC | .10 (.17) | .11 (.18) | .01 (.24) |
| Parietal | .12 (.17) | .13 (.18) | .01 (.23) |
| Striatum | .20 (.18) | .18 (.18) | -.02 (.24) |
| **T1w/T2w Ratio** |  |  |  |
| Left LPFC | .94 (.13) | .93 (.12) | -.01 (.03) |
| Right LPFC | .91 (.13) | .91 (.13) | -.001 (.04) |
| MPFC | .89 (.12) | .88 (.12) | -.01 (.04) |
| Parietal | .96 (.13) | .05 (.12) | -.01 (.04) |
| Striatum | 1.10 (.13) | 1.09 (.13) | -.01 (.03) |
| **2-back > 1-back Betas** |  |  |  |
| Left LPFC | 27.12 (18.10) | 21.60 (19.92) | -5.52 (28.27) |
| Right LPFC | 28.19 (19.69) | 20.60 (25.68) | -7.59 (31.89) |
| MPFC | 19.33 (15.51) | 15.21 (20.01) | -4.11 (24.22) |
| Parietal | 29.35 (21.33) | 27.24 (24.81) | -2.11 (28.18) |
| Striatum | 9.48 (8.85) | 4.76 (11.41) | **-4.72\* (14.01)** |

**Supplemental Figure 1.**
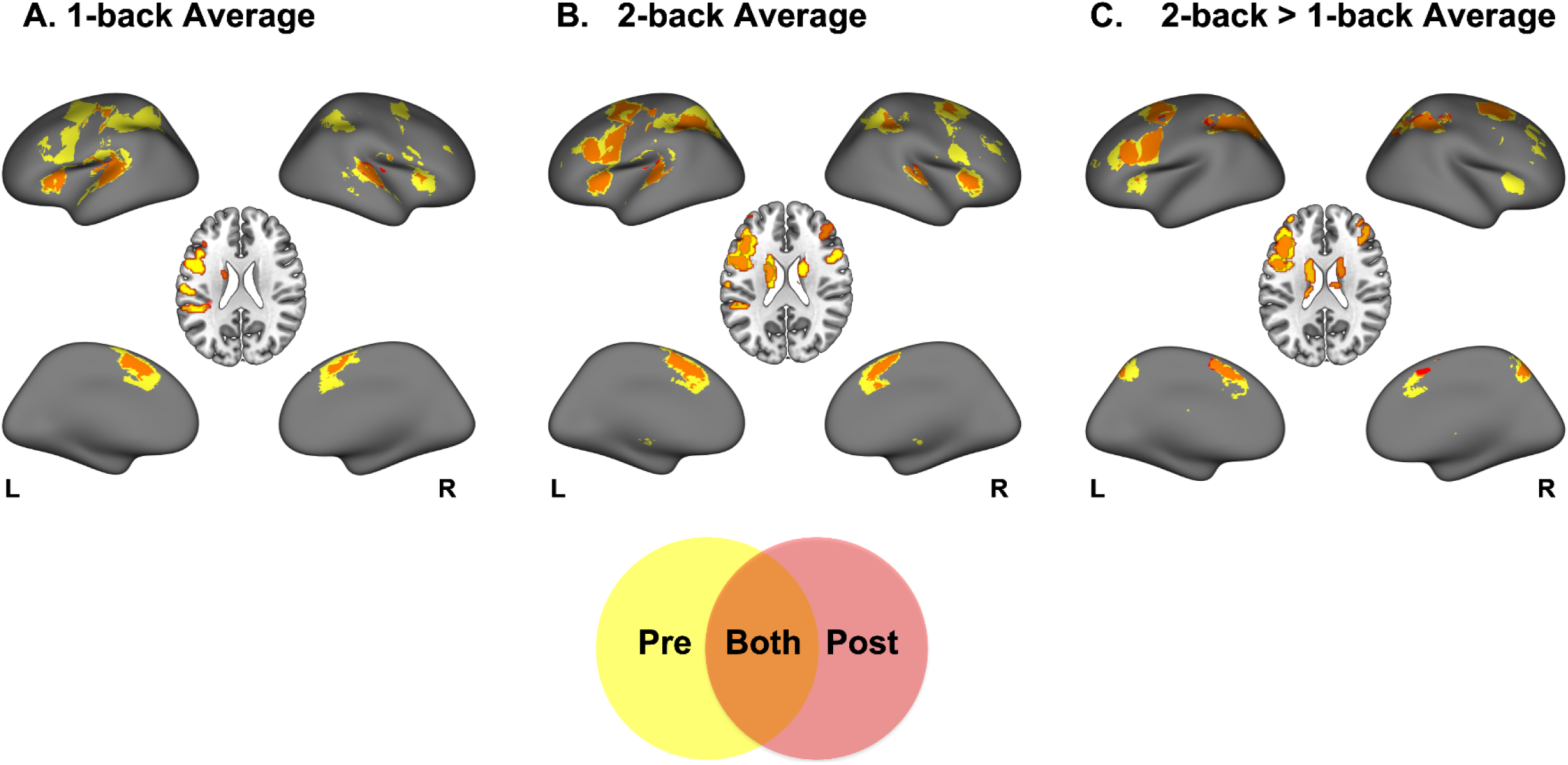
***N*-back task activation before and after adaptive *n*-back practice.** Average functional activation for (A) 1-back > baseline, (B) 2-back > baseline, and (C) 2-back > 1-back. Activation at baseline (pre-training) is shown in yellow, activation post-training is shown in red, and their overlap is shown in orange. Pre-training regions in the 2-back > 1-back contrast (shown in yellow in Panel C) were used as regions of interest in the VTA functional connectivity and T1w/T2w ratio analyses. Results are corrected for multiple comparisons at *z* = 4.0, *p* < 0.05. Axial slices are shown for *z* = 22.

**Supplemental Figure 2.**
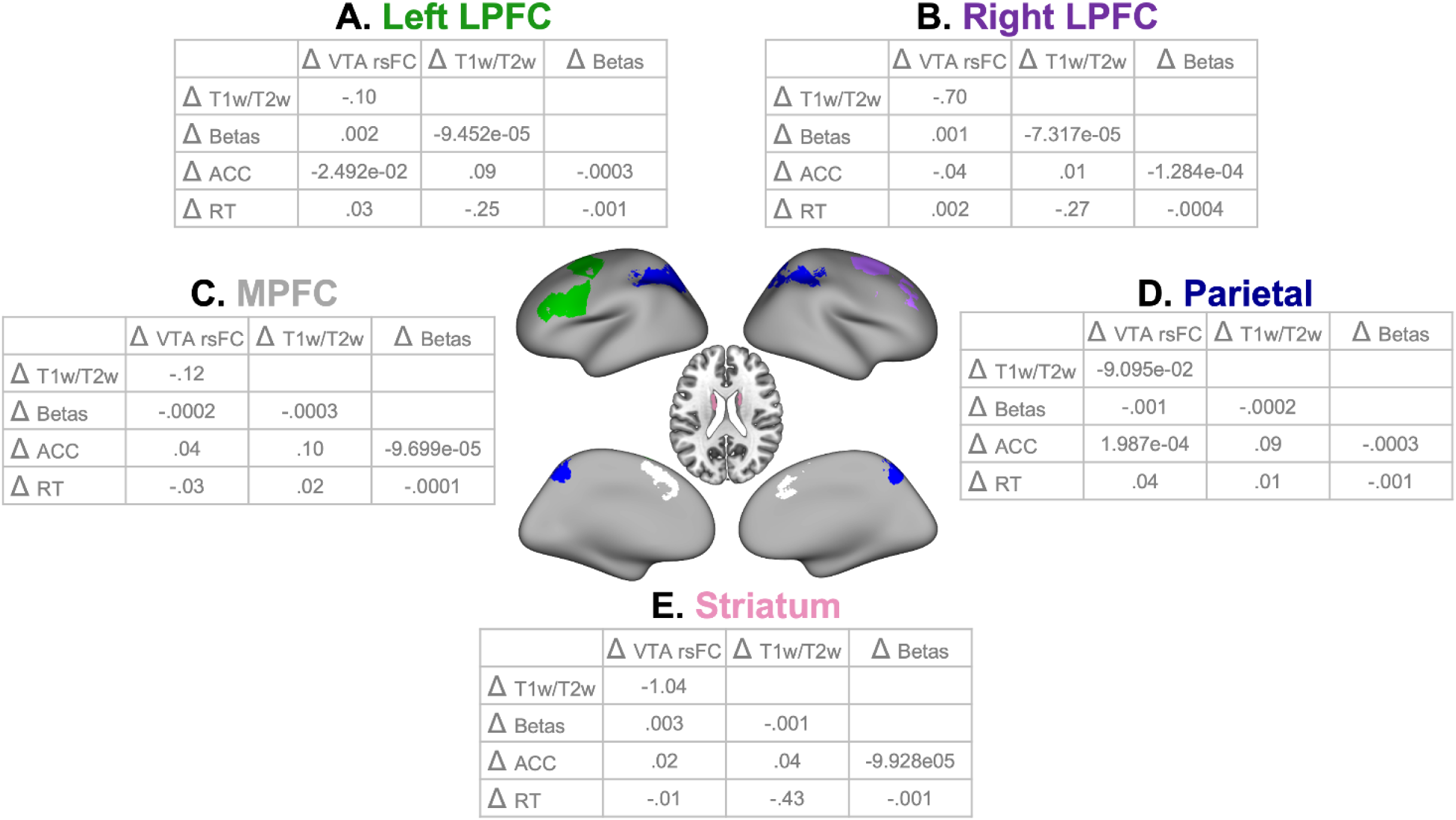
Beta coefficients for regression models examining associations between training-related changes in neural measures and working memory performance, rounded to the nearest hundredth. Axial slice visualized at *Z* = 22. Abbreviations: lateral prefrontal cortex (LPFC), medial prefrontal cortex (MPFC), resting-state functional connectivity between the ventral tegmental area and frontoparietal regions of interest (VTA rsFC), T1w/T2w ratio “myelin map” values (T1w/T2w), 2-back > 1-back beta values (Betas), *n*-back accuracy (ACC), *n*-back response times (RT). Statistical models control for age, sex, baseline working memory performance, baseline brain measures, and motion.

